# The study of hormonal metabolism of Trincadeira and Syrah cultivars indicates new roles of salicylic acid, jasmonates, ABA and IAA during grape ripening and upon infection with *Botrytis cinerea*

**DOI:** 10.1101/475053

**Authors:** João Coelho, Marilia Almeida-Trapp, Diana Pimentel, Flávio Soares, Pedro Reis, Cecília Rego, Axel Mithöfer, Ana Margarida Fortes

**Affiliations:** Universidade de Lisboa, Faculdade de Ciências de Lisboa, BioISI, Campo Grande, 1749-016, Lisboa, Portugal; Max Planck Institute for Chemical Ecology, D-07745 Jena, Germany; Instituto Superior de Agronomia, Universidade de Lisboa, Tapada da Ajuda, 1349-017 Lisboa, Portugal

**Keywords:** *Botrytis cinerea*, grape, phytohormones, necrotrophic pathogen, plant resistance, *Vitis vinifera*

## Abstract

Hormones play an important role in fruit ripening and in response to biotic stress. Nevertheless, analyses of hormonal profiling during plant development and defense are scarce. In this work, changes in hormonal metabolism in grapevine (*Vitis vinifera*) were compared between a susceptible (Trincadeira) and a tolerant (Syrah) variety during grape ripening and upon infection with *Botrytis cinerea.* Infection of grapes with the necrotrophic pathogen *Botrytis cinerea* leads to significant economic losses worldwide.

Peppercorn-sized fruits were infected in the field and mock-treated and infected berries were collected at green, *veraison* and harvest stages for hormone analysis and targeted qPCR analysis of genes involved in hormonal metabolism and signaling. Results indicate a substantial reprogramming of hormonal metabolism during grape ripening and in response to fungal attack. Syrah and Trincadeira presented differences in the metabolism of abscisic acid (ABA), indole-3-acetic acid (IAA) and jasmonates during grape ripening that may be connected to fruit quality. On the other hand, high basal levels of salicylic acid (SA), jasmonates and IAA at an early stage of ripening, together with activated SA, jasmonates and IAA signaling, likely enable a fast defense response leading to grape resistance/ tolerance towards *B. cinerea*.

The balance among the different phytohormones seems to depend on the ripening stage and on the intra-specific genetic background and may be fundamental in providing resistance or susceptibility. In addition, this study indicated the involvement of SA and IAA in defense against necrotrophic pathogens and gains insights into possible strategies for conventional breeding and/or gene editing aiming at improving grape quality and grape resistance against *Botrytis cinerea*.

## 1. Introduction

Hormones play a key role in the development and ripening of grapes and other fleshy fruits as well as in biotic stress response [1,2]. During grape ripening, significant physiological shifts occur including cell wall remodeling, accumulation of soluble sugars, aroma compounds and anthocyanins, decline of inducible host defense responses, among others. Most of these changes are thought to be regulated by a complex interplay of hormonal signals involving ethylene, ABA, brassinosteroids, jasmonates, polyamines, cytokinins and auxins [1,3]. Interestingly, similar phytohormones are regulated in the host in response to pathogens, and this regulation is modulated during fruit ripening when fruits show increased susceptibility to pathogens [2,4].

Fleshy fruits can be classified into two groups, climacteric and non-climacteric. Climacteric fruits such as tomato show a concomitant increase in respiration and ethylene biosynthesis upon initiation of ripening. In non-climacteric fruits such as grapes the respiratory burst and rise in ethylene production are absent at the onset of ripening (reviewed by Fortes *et al.*, 2015) [1]. No single master switch controlling ripening initiation, such as ethylene in climacteric fruits, has yet been uncovered for non-climacteric fruits [5].

Abscisic acid (ABA), brassinosteroids (BRs), and ethylene have been suggested to promote ripening through complex interactions, while auxin delays some ripening associated processes and also interacts with other phytohormones such as ABA and ethylene [6]. In particular, in non-climacteric grape fruits ABA has been proposed as the main signal triggering the onset of ripening-associated processes since it peaks at *veraison*, along with initiation of berry softening and skin coloration [6,7]. Increases in the levels of ABA also influence the accumulation of sugars and their enhanced uptake and storage can be stimulated by exogenous application of ABA before *veraison* [8]. Treating grape berries with ABA at pre-*veraison* or *veraison* can also contribute to improve anthocyanin levels and the biosynthesis of phenylpropanoids [9,10]. The endogenous ABA content is determined by the dynamic balance between biosynthesis, conjugation and catabolism [11].

Auxins have an important role in fruit growth, but act as inhibitors of ripening in both climacteric and non-climacteric fruits [12]. Low auxin levels seem to be required at the onset of grape ripening, which is associated with an increase in the conjugated forms of indole-3-acetic acid [13]. Auxin delays increase in berry size, sugar accumulation, and anthocyanin content; in fact, auxin negatively regulates ABA-induced ripening processes [14–16]. Nevertheless, a core set of several genes involved in auxin signaling has been found to be differentially expressed during grape ripening of three cultivars [17]. In tomato, altered *AUXIN RESPONSE FACTOR 2* expression leads to modified contents in abscisic acid, cytokinins and salicylic acid highlighting that auxin signaling intersects hormonal signals in the regulation of fruit ripening [18].

As it occurs for auxins, jasmonic acid (JA) levels are higher during early development and decrease during ripening, which appears to enable the onset of this stage. Furthermore, genes involved in jasmonate biosynthesis were found to be less expressed during and after *veraison* [19]. Nevertheless, jasmonic acid has been suggested to be involved in ripening by influencing the coloring, softening and aroma of the fruit [20]. A volatile form of JA, methyl jasmonate, is capable to promote red coloring, increase in anthocyanin content and promotion of volatiles’ synthesis when exogenously applied on fruit [21,22]. Additionally, JA is known to regulate plant immune responses in particular against necrotrophic pathogens, e.g. *Botrytis cinerea* [23,24]. *B. cinerea* is a fungus to which several varieties of grapevine are highly susceptible in particular the Portuguese Trincadeira [4].

Salicylic acid has been described to be involved in the activation of plant defenses against pathogenic fungi, biotrophs and hemibiotrophs, but it may enhance susceptibility to necrotrophs by antagonizing the JA signaling pathway and by inhibition of auxin signaling [23,25,26]. Salicylic acid signaling was shown to have an important role in necrotrophic colonization stage of *Colletotrichum* on tomato fruit by inducing cell death and likely as a means to suppress JA mediated defense response [27]. However, in unripe tomato fruit the transgenic NahG tomato line, which does not accumulate SA, showed susceptibility to *Botrytis* [28] which suggests that SA might contribute to resistance of unripe fruits to *Botrytis*.

Besides SA and jasmonates, which are conserved positive regulators of plant defense, auxins are hormones with emerging roles in plant defense response [29–31]. Previous studies suggested that they may also be involved in the grape response to *Botrytis* attack since in the transcriptome analysis of infected grapes an enrichment in the functional class of auxin signaling genes was noticed [4]. Interestingly, an interaction between auxin and jasmonic acid was previously shown to occur in resistance to necrotrophic pathogens [32]. On the other hand, ABA has been described as a hormone that can either induce or repress plant defense depending on the specific plant–pathogen interaction [26,33]. It is generally considered that ABA suppresses plant resistance mechanisms by antagonizing SA- and JA/ET-dependent immune responses [34,35], though few examples associated ABA with disease resistance [33]. Additionally, during infection, certain plant pathogens can directly produce ABA or induce its synthesis in the host in order to accelerate fruit ripening and therefore fruit susceptibility [36]. Overall synthesis and perception of different stress hormones, along with their relative contents and interactions, seem to be crucial for plant resistance against pathogens.

Previously, transcriptome analysis of *Vitis vinifera* cv. Trincadeira berries upon infection with *B. cinerea* suggested the putative involvement in grape defense of jasmonic acid, ethylene, and auxins [4]. This innovative study focused on the fruit organ due to the fact that defense mechanisms against necrotrophic, biotrophic, or hemibiotrophic pathogens have been documented mostly for vegetative tissues [26]. In the present study, were compared a highly susceptible variety (Trincadeira) and a highly resistant/ tolerant variety (Syrah) of grape [4,37]. New developmental stages were taken into account allowing deepening the understanding of hormone regulation during grape ripening and grape defense against *Botrytis cinerea.* The expression of genes related to hormonal metabolism and signaling, as well as the analysis of the hormonal profile was carried out in order to unravel cultivar specific mechanisms involved in the onset of ripening and in effective defense.

## 2. Materials and methods

### 2.1 Infection of berries and sample collection

Field experiments were conducted in an experimental vineyard with 15-year-old grapevines (rootstock '140Ru') at the Instituto Superior de Agronomia, University of Lisbon, Portugal. Infections of *Vitis vinifera* cv Trincadeira and Syrah berries with *Botrytis cinerea* were made in early June of 2016. The *B. cinerea* isolate used was obtained from diseased grapevine plants and maintained in potato dextrose agar (Difco, Detroit, MI, USA), at 5 °C. Conidia production was achieved by exposing inoculated Petri dishes with potato dextrose agar to continuous fluorescent light, at 24 °C. Conidia were harvested from 14- to 20-day-old cultures and collected by rubbing with phosphate buffer (0.03 M KH_2_PO_4_), filtered through cheesecloth to remove mycelia, and the concentration adjusted to 10^5^ conidia ml^-1^. The infections were made by spraying berry clusters with a conidial suspension at the developmental stage of peppercorn size (stage EL29), following the procedure by Agudelo-Romero *et al.* [4]. Collection of Trincadeira and Syrah samples after visual inspection of symptoms was performed at three different stages of development/ ripening: green (stage EL32), *veraison* (stage EL35) and harvest (stage EL38) according to the modified E–L system [38]. Five to six replicates were obtained for each stage of development and for each treatment (control sprayed with phosphate buffer), and all berry clusters were briefly transported in ice to the laboratory, frozen in liquid nitrogen and kept at −80 °C until further use. Previous phenotypic, molecular and biochemical analyses showed that treatment of grapes with buffers had no visible impact on grape ripening [39]. Prior to extraction for transcriptional and metabolic analyses, the seeds were removed. Four to six biological replicates were used for quantification of hormones and some of these samples were pooled in order to yield three independent biological replicates for study of gene expression.

### 2.2 *DNA extraction and assessment of* Botrytis cinerea *infection with qPCR*

DNA extraction was performed according to Lodhi *et al.* [40] with modifications: a 2 M KAc treatment (1 h on ice) to precipitate polysaccharides was added to the protocol and performed before RNase A treatment. Fungal biomass accumulation and infection level was determined by qPCR amplification of the *B. cinerea* polygalacturonase 1 (PG1). DNA accumulation levels were linearized with the formula 2 ^-(BcPG1^ ^Ct-VvACT^ ^Ct)^ using the grape actin gene as reference.

### 2.3 RNA extraction

RNA extraction was carried out according to Fortes *et al.* [19] with slight modifications: the KAc treatment for polysaccharides removal was done before overnight RNA precipitation with LiCl in order to avoid an additional precipitation with sodium acetate and ethanol. A DNase treatment was performed according to the supplier’s instructions (Invitrogen, San Diego, CA, USA). RNA was further purified using the Spectrum™ Plant Total RNA Kit (Sigma-Aldrich).

### 2.4 Real-time PCR

First-strand cDNA was synthesized from 2 μg of total RNA as described previously [19]. Real-time PCRs were performed using the StepOneTM Real-Time PCR System (Applied Biosystems, Foster City, CA, USA). Cycling conditions were 95 °C for 10 min, followed by 42 cycles of 95 °C for 15 s and 56 ºC- 60 °C for 40 s. Relative expression data were derived from three biological replicates and duplicate technical replicates (performed in separate plates). The standard curve was built using a serial dilution of mixtures of all cDNAs analyzed. Primer efficiencies (in between 85 and 105%) were calculated using 4- fold cDNA dilutions (1:1, 1:4, 1:16, 1:64, and 1:256) in triplicate as well as checking for amplification in a negative control without DNA. In order to obtain relative expression of the genes under study, data were normalized using the expression curves of the actin gene (VIT_04s0044g00580) and elongation factor 1α gene (VIT_06s0004g03220). These genes are the most stable according to NormFinder software [4]. Selection of the genes involved in hormonal metabolism for qPCR analysis was based on previous microarray data [4], analyses of hormonal pathways (Supplementary Figure S1) and bibliography. All primers used are shown in Supplementary Table S1.

### 2.5 Hormonome analysis

Stock solutions of each original phytohormone standard were prepared at 1 mg ml^-1^ in MeOH. For deuterated compounds, stock solutions were prepared in acetonitrile at 100 μg ml^-1^. Working solutions of original phytohormones standards were prepared by diluting stock solutions in MeOH:water (7:3), at different concentration for each phytohormone depending on the range of the calibration curve: ABA and IAA at 100 μg ml^-1^; JA and SA at 200 μg ml^-1^; OPDA at 50 μg ml^-1^; and JA-Ile at 40 μg ml^-1^. The internal standard stock solutions (d5-JA, d6-ABA, d4-SA, and d5-IAA) were combined and diluted in MeOH:water (7:3) ratio, resulting in the extraction solution. The final concentrations were 10 ng ml^-1^ for both d4-SA and d5-IAA, and 20 ng ml^-1^ for both d5-JA and d6-ABA.

The samples (5 to 6 biological replicates) were freeze dried at −40 °C for three days. One ml of extraction solution containing the internal standards (d5-JA, d6-ABA, d5-IAA, and d4-SA), prepared as described previously, was directly added. The samples were briefly mixed with a vortex, and spiked with phytohormones standards as described in Almeida Trapp *et al.* [41]. The spiked samples were shaken for 30 min and centrifuged at 16,000 x *g* and 4 °C for 5 min. The supernatant was transferred into a new micro-centrifuge tube and dried in speed vac. After drying, 100 μl of MeOH were added to each sample, which were then mixed with a vortex and centrifuged at 16,000 x *g* and 4 °C for 10 min. The supernatant was analyzed by HPLC-MS/MS (high performance liquid chromatography-mass spectrometry) as described [41].

### 2.6 Statistical analysis

In order to identify the significance of genotype or treatment on gene expression or phytohormones content, all data were analyzed by two-way ANOVA. In the cases where the residuals were not normally distributed and/or the variance was not equal among the groups, the data were prior transformed, to be further tested with ANOVA. For the analyses of phytohormones, some outliers were identified by two-tailed Dixon’s Q-Test (with Q = 95%) and excluded before the two-way ANOVA. All data were analyzed with Sigma Plot (version 12.0).

### 2.7 Anthocyanin quantification

Anthocyanin concentration was measured as described previously [19]. Grapes were frozen in liquid nitrogen, seeds removed, freeze-dried for 72-96 h at – 40 °C and then 100 mg of the powder extracted in 1.5 ml TFA (Trifluoroacetic acid)/methanol/H_2_O (0.05/80/20, v/v/v). Samples were vortexed for 1 min and then anthocyanins were extracted for 1 h on ice in Eppendorf tubes. The mixture was then centrifuged for 30 min at 13,000 rpm at 4 °C. A 50 μl or 100 μl of this sample was diluted to 1 ml in extraction solution for *veraison* and harvest samples. The solution was mixed and allowed to sit for 5 min before reading the absorbance at A_520_. Total relative anthocyanin concentration was expressed as the absorbance value at 520 nm g^−1^ of freeze-dried weight.

## 3. Results

### 3.1 *Phenotypic characterization of infected and mock-treated grape berries and assessment of* Botrytis cinerea *infection*

Berries were inoculated with *B. cinerea* conidial suspension at the peppercorn-sized fruits stage corresponding to developmental stage EL29 according to the Coombe (1995) classification [38]. The ripening stages investigated in this study were identified as EL32 characterized by hard green berries, EL35 corresponding to *veraison* when anthocyanin accumulation initiates and EL38 corresponding to fully ripe berries (harvest stage).

Figure 1 shows infected Trincadeira and Syrah clusters corresponding to EL32, EL35 and EL38. Trincadeira clusters presented already at EL32 a high infection level with *B. cinerea*. Evaluation of sample infection was performed by visual inspection and additionally by qPCR using primers specific to the fungal genomic DNA (Fig. 2). Syrah present very mild symptoms at EL32 which was corroborated by the qPCR analysis (Fig. 2). Though symptoms increase during ripening Syrah still presented at EL38 a lower level of infection than Trincadeira which is not surprising since Syrah is known to be tolerant/ resistant to the disease under normal in-field conditions [37].

**Figure 1:**
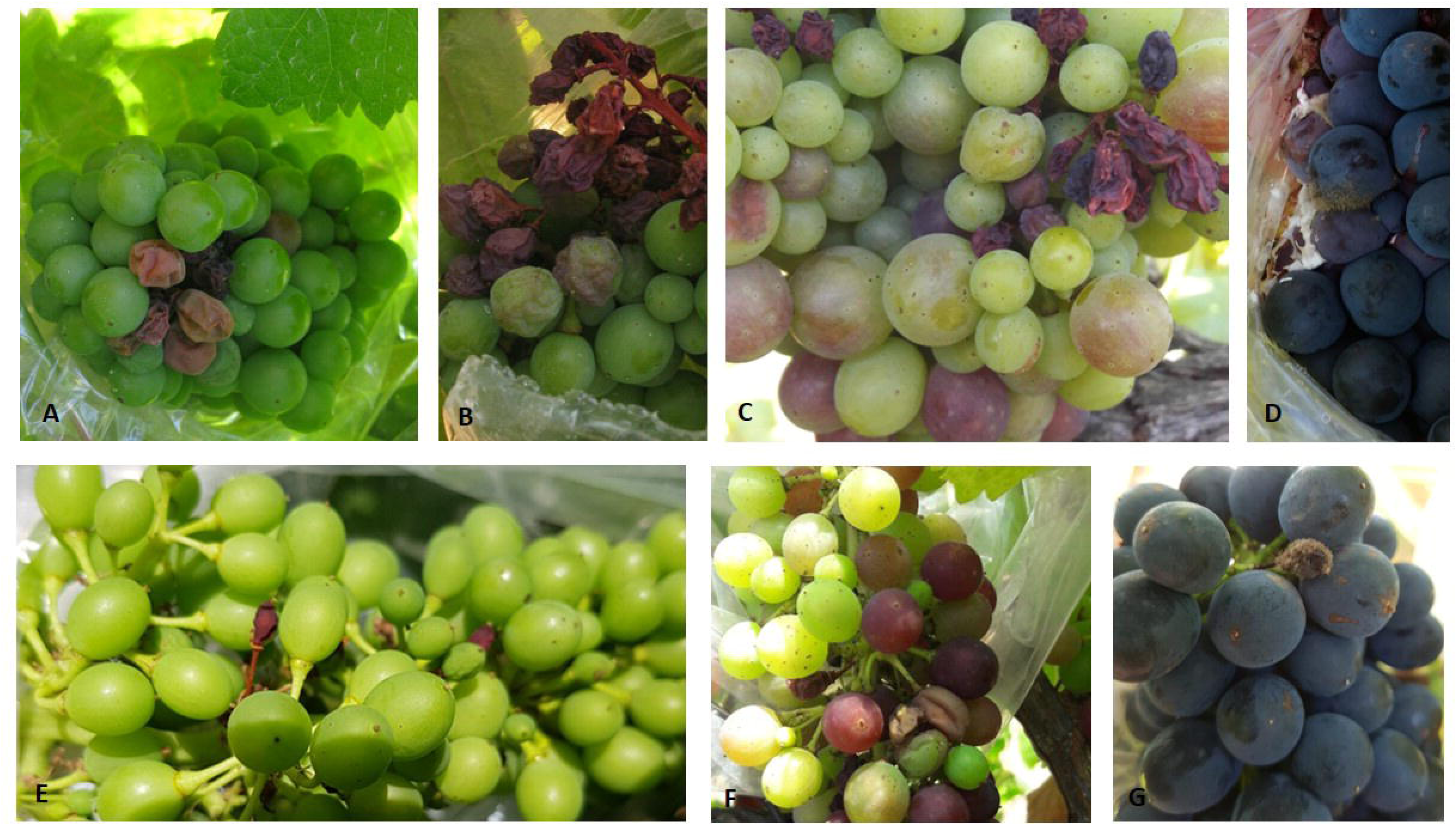
– Clusters of Trincadeira and Syrah grapes infected with *B. cinerea.* A, B Clusters of Trincadeira at EL32. C, Cluster of Trincadeira at EL35. D, Cluster of Trincadeira grapes at EL38. Sporulation of the fungus is observed in some berries of the cluster. E, Cluster of Syrah at EL32 with no significant symptoms. F, Cluster of Syrah at EL35. G, Cluster of Syrah grapes at EL38.

**Figure 2:**
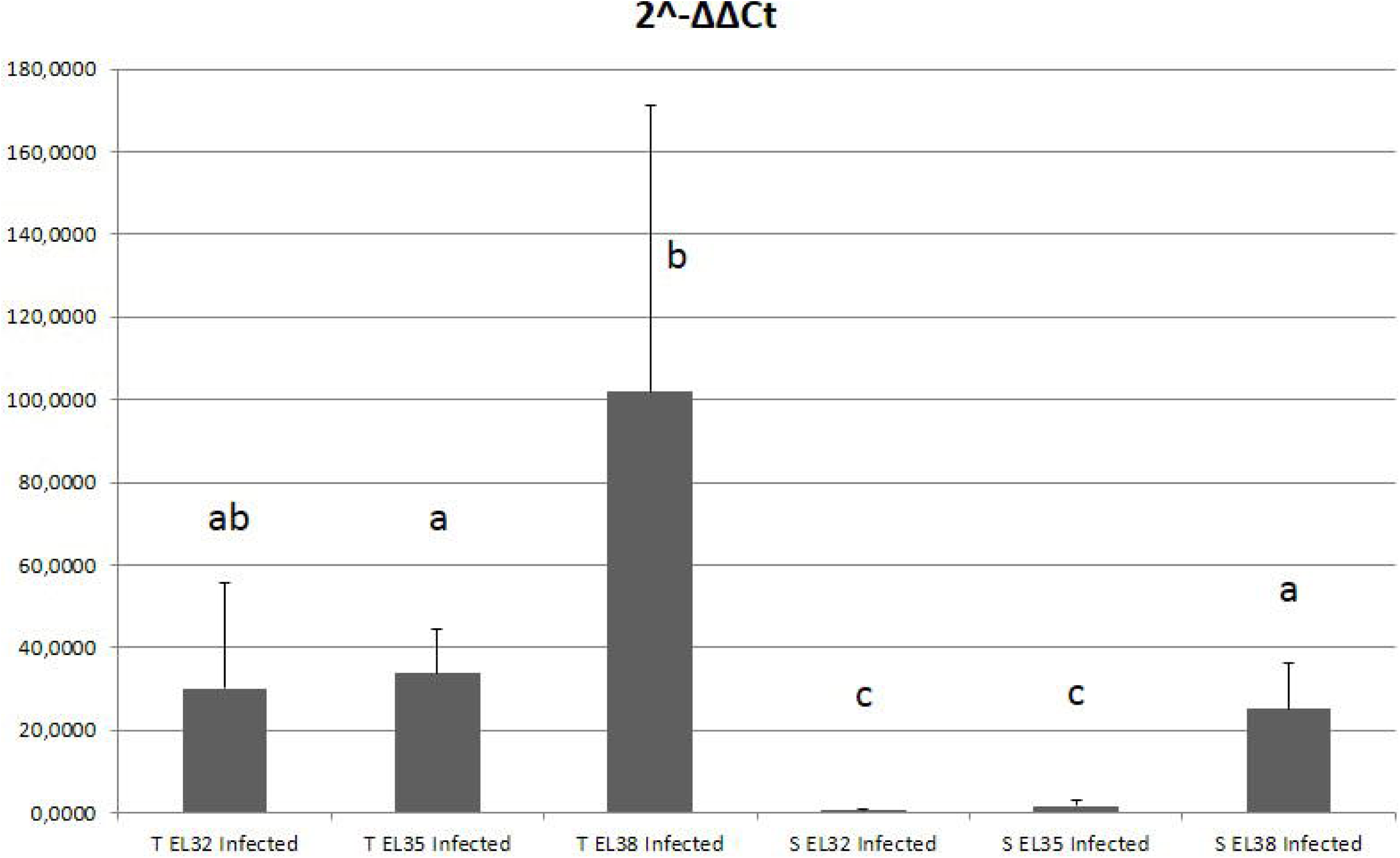
Evaluation of infection level by qPCR using primers specific to the fungal genomic DNA (*BcPG1*). Syrah presented very mild infection at EL32 whereas Trincadeira was severely infected. Bars represent the standard deviation error. Data were analyzed with Welch's t-test. Different infection level is given by different lettering (p< 0.05).

The assessment of ripening stages took into account the fresh berry weight and the content in anthocyanins present in healthy berries (Fig. 3). Content in anthocyanins has been described as a reliable indicator of the ripening stage [42]. Additionally, recent metabolomics data did not show significant differences in the main ripening parameters between control berries of the two cultivars (data not shown). As results of *B. cinerea* infection on Trincadeira, a decrease in berry weight in infected compared with non-infected samples and an increase in anthocyanin content at EL35 were detected, indicating that infected berries were riper than non-infected berries as previously noticed and supported by molecular and metabolic data [4]. The acceleration of ripening in Trincadeira berries induced by *B. cinerea* constitutes a strategy of the fungus to lead to increased susceptibility of the fruit by inhibiting defenses that are active in the green fruit [4]. The acceleration of ripening as assessed by berry weight and anthocyanin content was not observed for less infected Syrah berries.

**Figure 3:**
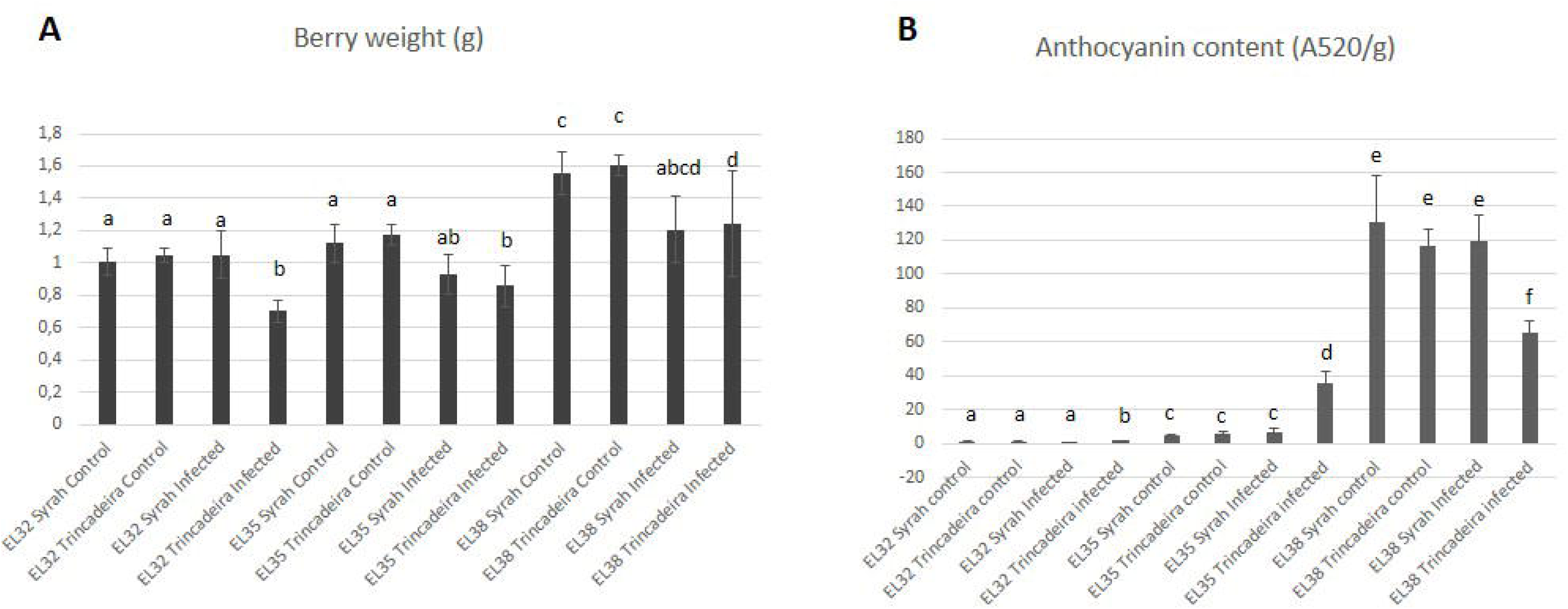
Phenotypic characterization of infected and mock-treated grape berries. (A, B) Fresh berry weight (A; g) and total anthocyanin content (B; absorbance at 520 nm g^-1^ of freeze-dried material) of infected and control berries at developmental stages of EL32, EL35 and EL38. Bars represent the standard deviation error. Data were analyzed with Welch's t-test. Different infection level is given by different lettering (p< 0.05).

On the other hand, the content in anthocyanins at EL38 decreased significantly in the severely infected Trincadeira but not in Syrah. This decrease in anthocyanins was previously observed in berries severely infected with powdery mildew [43] and in grape skins of *Botrytis*-affected berries [44].

### 3.2 Hormonal metabolism of Trincadeira and Syrah cultivars during grape ripening

For jasmonate analyses we focused on the bioactive form, jasmonoyl-isoleucine conjugate (JA-Ile) [45], and its biosynthetic precursor 12-*oxo*-phytodienoic acid (OPDA). At EL32, both cultivars showed similar amounts of OPDA and started decreasing from the onset of grape ripening until EL38 (Fig. 4). In case of JA-Ile, a decrease from EL32 to EL38 was also found in both cultivars; however, at EL32 the JA-Ile concentration in Syrah grapes was about three times higher than in Trincadeira and at EL35 it was still higher. The expression of a gene coding for allene oxide synthase (AOS) involved in OPDA synthesis (Supplementary Figure S1) showed a pronounced increase during ripening of Trincadeira grapes. Another biosynthetic gene encoding 12-oxophytodienoate reductase 1 (OPR1) and further involved in JA-Ile biosynthesis tended to be slightly higher at the green stage (EL32) in both cultivars in accordance with the concentration of JA-Ile. However, at EL35 Trincadeira presented higher expression of this gene but lower JA-Ile levels. A gene coding for JAZ8, a repressor of jasmonate signaling [46] was up-regulated at EL38 in both Trincadeira and Syrah grapes. *MYC2* expression involved in jasmonate-dependent transcriptional activation [31], presented an expression pattern during grape ripening similar to *JAZ8* but with a lower fold-change level. Previous microarray analysis showed that mRNAs involved in the biosynthesis of jasmonates, namely those coding for OPR1 were less abundant at EL35 and EL36 in Trincadeira grapes [19]. Nevertheless, a gene coding for an allene oxide synthase (AOS) involved in jasmonate biosynthesis was strongly up-regulated at EL35 and EL36 in this previous study. In the present analyses, both cultivars collected in a different *terroir* from previously one [19] presented an increase in the expression of a gene coding for a specific isoenzyme of AOS during grape ripening.

**Figure 4:**
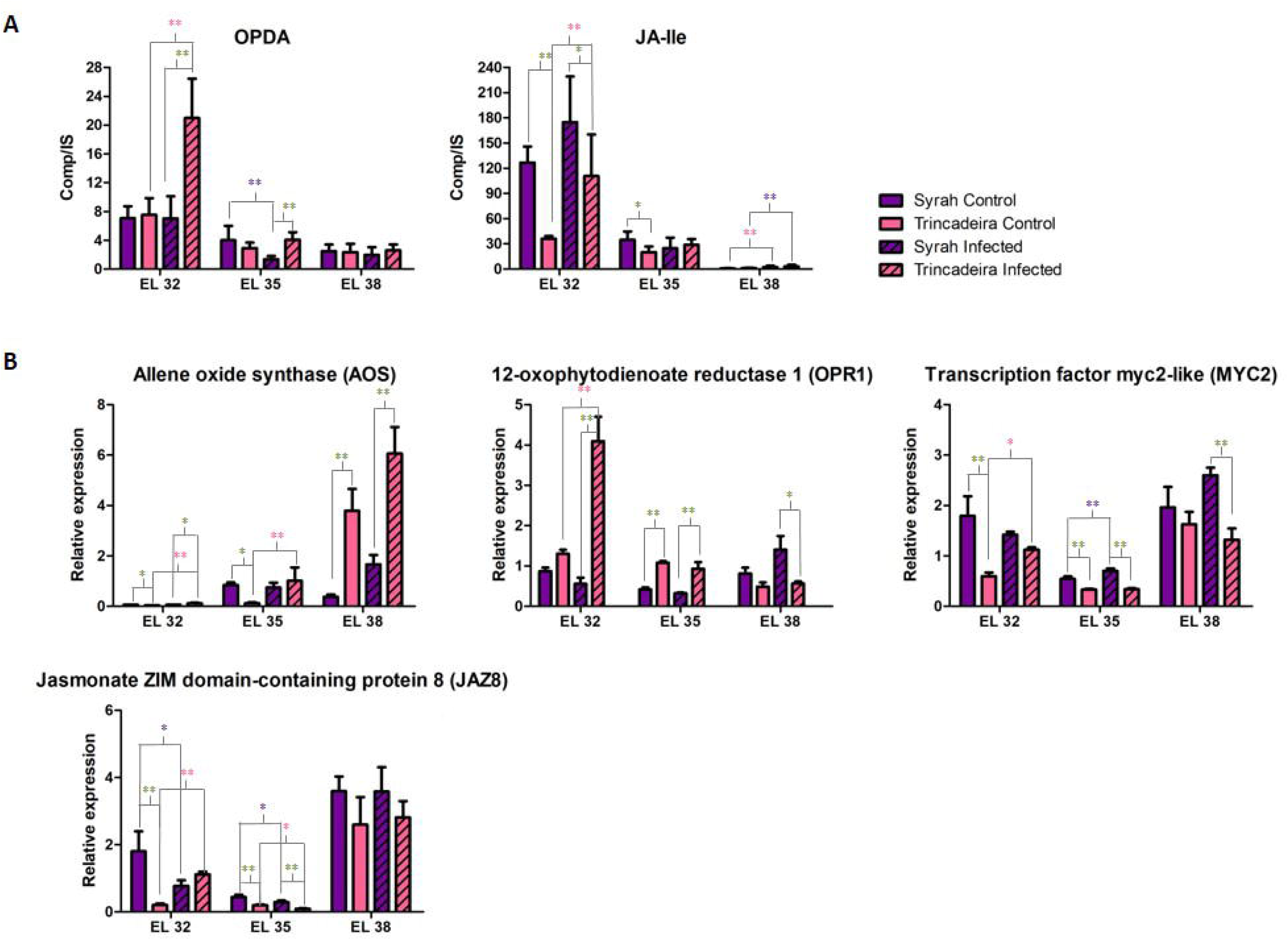
Metabolism of jasmonates in Trincadeira and Syrah grapes at three development stages: green (EL32), *veraison* (EL35) and harvest (EL38). A, Hormonal quantification of jasmonates (OPDA and JA-Ile); B, Expression of genes involved in jasmonates metabolism: *AOS*, *OPR1, MYC2* and *JAZ8*. Data were analyzed by two-way ANOVA: * P<0.05; **P<0.01. Different colors present either intra or inter-variety comparisons.

Similar to jasmonates the content in IAA showed a tendency to decrease in Syrah and Trincadeira grapes during ripening (Fig. 5). The highest expression levels of the gene coding for Indole-3-acetic acid-amido synthetase, GH3.2, involved in auxin inactivation by conjugation [47] were at stage EL32 in accordance with IAA content. The expression of this gene lowers along grape ripening, and more abruptly in Syrah grapes. Expression of *IAA-amino acid hydrolase 6* was higher at EL32 compared with EL35 but tended to increase at EL38. Altogether, these results suggest that free IAA levels are regulated at green stage mainly by conjugation and at harvest stage mainly by hydrolysis of conjugates. In both cultivars, the amount of IAA reached its lowest content at the harvest stage, though higher in Trincadeira.

**Figure 5:**
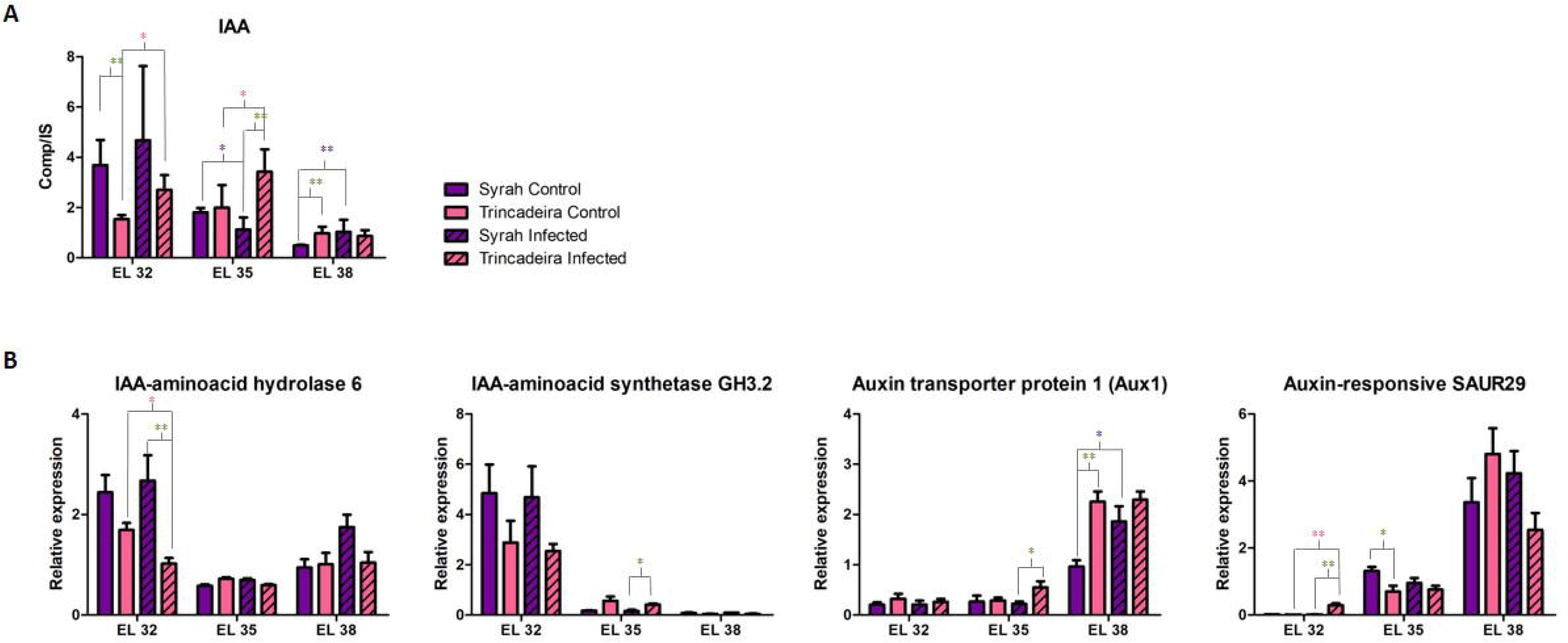
Metabolism of auxins in Trincadeira and Syrah grapes at three development stages: green (EL32), *veraison* (EL35) and harvest (EL38). A, Hormonal quantification of Indole-3-acetic acid (IAA); B, Expression of genes involved in auxins’ metabolism: *IAA hydrolas*e *8*, *indole-3-acetic acid-amido synthetase, GH3.2*, *Aux1* and *Auxin responsive SAUR29*. Data were analyzed by two-way ANOVA: * P<0.05; **P<0.01. Different colors present either intra or inter-variety comparisons.

Small Auxin Up RNAs (SAURs) are the largest family of early auxin response genes (Supplementary Figure S1). Members of the auxin-responsive SAUR gene family can act as negative regulators of auxin biosynthesis [48]. The *auxin-responsive SAUR29* gene presented lower levels of expression at stage EL32 in accordance with the peak of auxin content observed in both cultivars at this stage. As the expression of the *auxin-responsive SAUR29* increased, auxin concentration decreases, until the gene reached its peak of expression at stage EL38. The same was verified for *Aux1* involved in auxin transport [49]. Interestingly, the expression of *Aux1* was higher in Trincadeira at EL38 whereas the expression of *auxin-responsive SAUR29* was higher in Syrah at EL35.

Regarding SA concentrations they steadily decreased across the various stages of ripening, both in Syrah and in Trincadeira grapes (Fig. 6). The expression of genes coding for enhanced disease susceptibility (EDS1) and phytoalexin deficient 4 (PAD4) involved in SA signaling [50] also followed this trend in both cultivars; the same observation was obtained for Trincadeira collected in another *terroir* and season [19]. Out of the two cultivars, Syrah displayed at EL32 higher basal levels of SA than Trincadeira and higher expression of *PAD4*.

**Figure 6:**
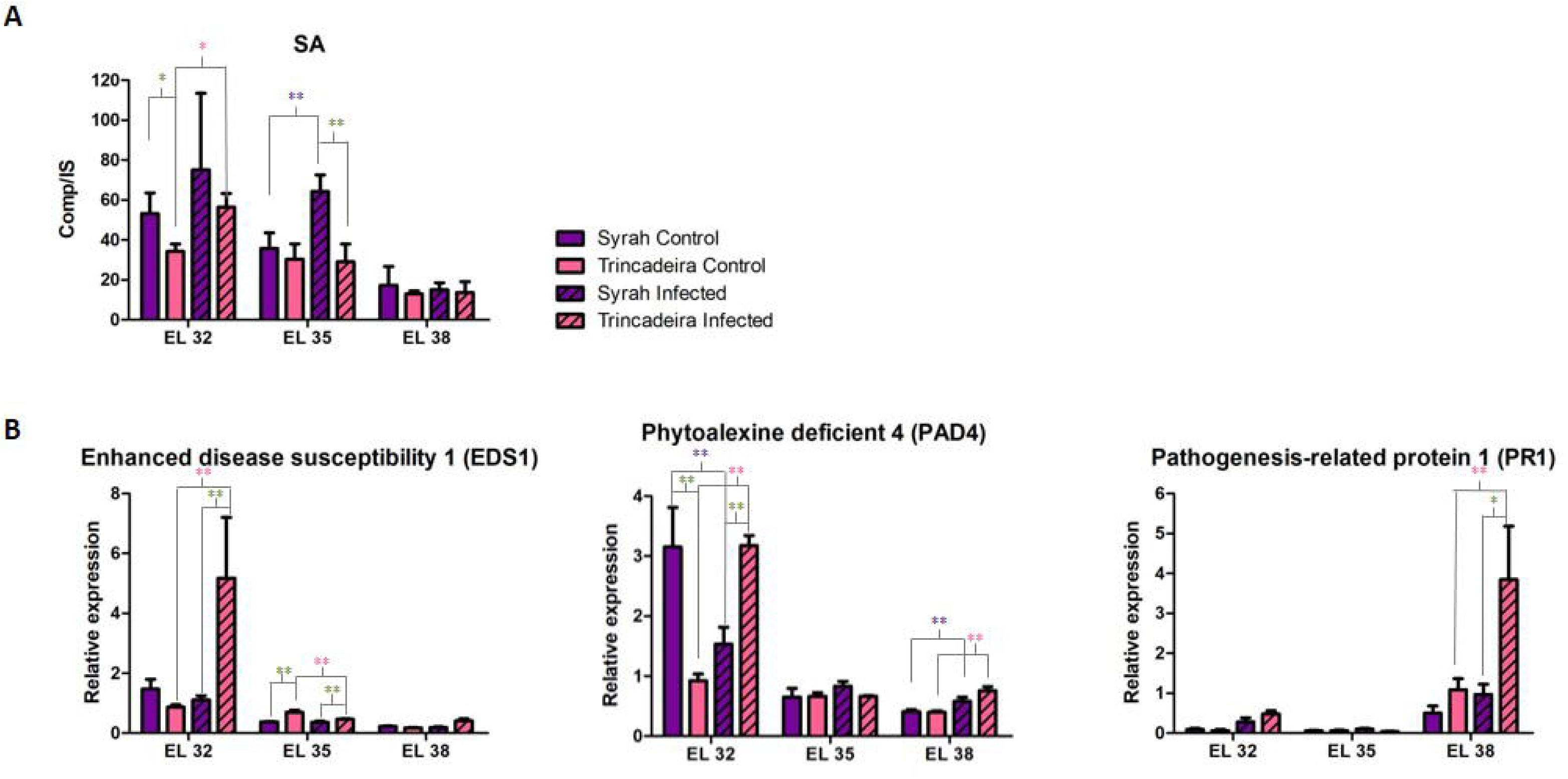
Metabolism of SA in Trincadeira and Syrah grapes at three development stages: green (EL32), *veraison* (EL35) and harvest (EL38). A, Hormonal quantification of SA; Expression of genes involved in SA signaling: *EDS1*, *PAD4* and *PR1*. Data were analyzed by two-way ANOVA: * P<0.05; **P<0.01. Different colors present either intra or interspecies comparisons. Different colors present either intra or inter-variety comparisons.

While jasmonates and auxins decreased during ripening, ABA concentration increased drastically at *veraison* in both Trincadeira and Syrah grapes and decreased again at harvest (Fig. 7). The levels of expression of gene coding for 9-cis-epoxycarotenoid dioxygenase (NCED) involved in ABA biosynthesis (Supplementary Figure S1) was higher at *veraison* in both cultivars and about two times in Syrah at EL35. These data are in general in accordance with the variations in ABA content showed by hormone quantification. Interestingly, the gene coding for ABA receptor PYL4 RCAR10 involved in ABA-mediated signaling pathway [51] was more expressed during the green stage, especially in Syrah grapes. In fact, at EL32 when compared with Trincadeira, Syrah grapes presented higher expression of *NCED*, and *ABA receptor PYL4 RCAR10*, though levels of ABA were similar for both cultivars. The expression of this ABA receptor was more noticeable in Trincadeira grapes at harvest, possibly related to a higher content in ABA in this cultivar at this stage. Differences in ABA content in the two cultivars during grape ripening seem to be at least partially regulated by ABA catabolism involving ABA 8'-hydroxylase since higher expression of the gene coding for this enzyme was noticed for Syrah at EL35 and EL38.

**Figure 7:**
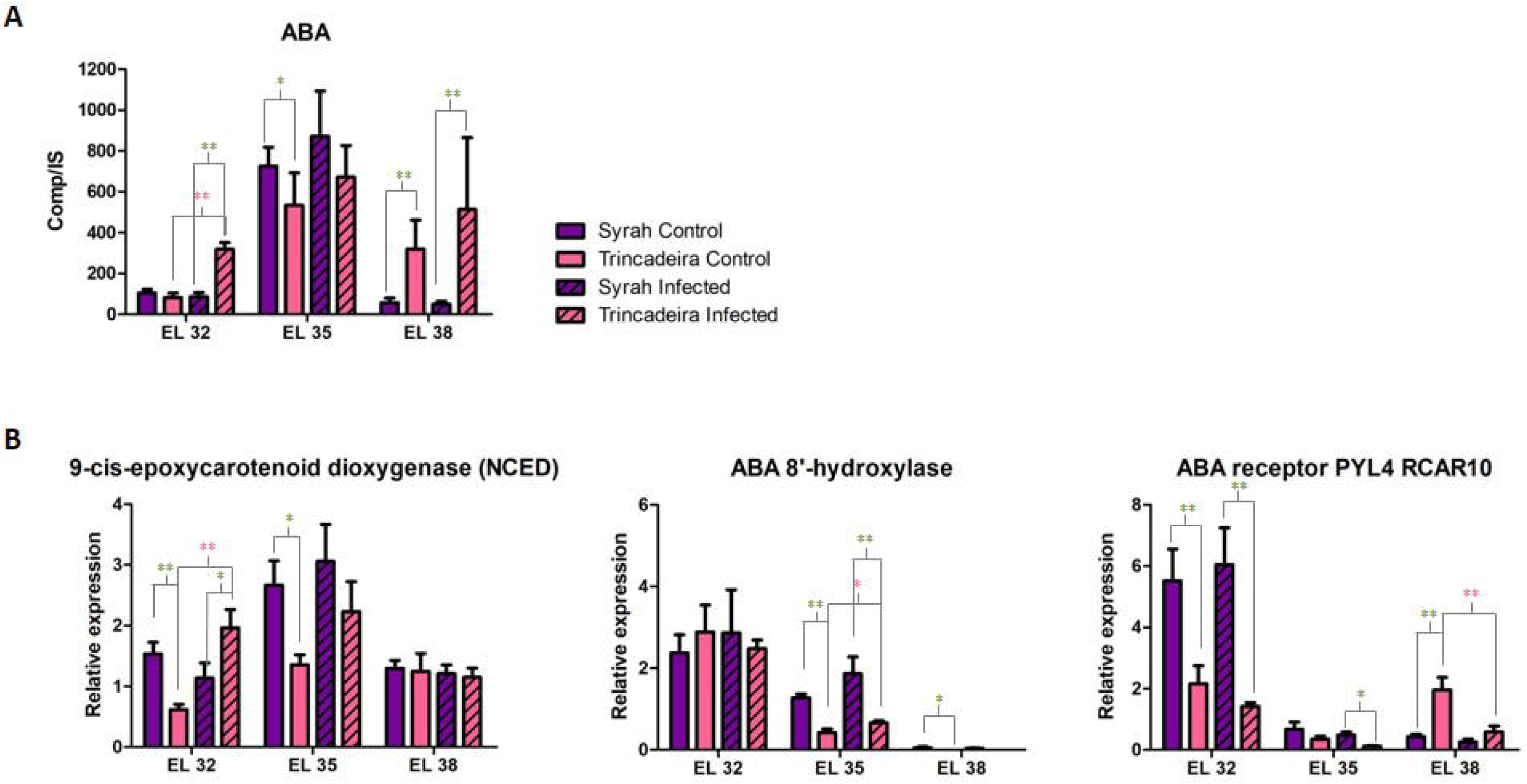
Metabolism of ABA in Trincadeira and Syrah grapes at three development stages: green (EL32), *veraison* (EL35) and harvest (EL38). A, Hormonal quantification of ABA; Expression of genes involved in ABA metabolism: *9-cis-epoxycarotenoid dioxygenase (NCED)*, *ABA receptor PYL4 RCAR10*, *ABA 8’-hydroxylas*e. Data were analyzed by two-way ANOVA: * P<0.05; **P<0.01. Different colors present either intra or interspecies comparisons. Different colors present either intra or inter-variety comparisons.

### 3.3 *Hormonal metabolism of Trincadeira and Syrah grapes upon infection with* Botrytis cinerea

Concerning jasmonates, at EL32, the amount of OPDA in Syrah grapes was maintained after infection, while in the susceptible Trincadeira variety, which already showed heavy symptoms, OPDA level increased (Fig. 4). The same held true for JA-Ile. However, Syrah grapes displayed higher basal levels of JA-Ile comparing to Trincadeira along with higher expression of *MYC2* and *JAZ8*. Regarding the expression of genes involved in the synthesis of jasmonic acid, it was noted that at green stage expression of *AOS* was very low in infected samples of both cultivars though differences could be noted between them. However, at harvest expression of *AOS* increased in infected berries of Syrah and Trincadeira, being highly expressed in the later one. The expression of this gene is in general in agreement with the content in JA-Ile at harvest stage. On the other hand, *OPR1* expression peaked at the green stage, which is in accordance with the concentration of JA-Ile. Particularly, it illustrates the increase in JA-Ile concentration in Trincadeira grapes upon infection accompanied by higher expression of *JAZ8* whereas the opposite was noticed in Syrah regarding the expression of this repressor of jasmonic acid signaling. *OPR1* expression decreased at *veraison* accompanied by a decrease in the content in JA-Ile. Interestingly, at EL38 infected Syrah grapes seem to present higher expression of *OPR1* than Trincadeira but this was not reflected in increased JA-Ile levels. It should be noted that only at this stage Syrah grapes presented heavy symptoms of infection.

Regarding auxins, the content in IAA increased in infected Trincadeira grapes at EL32 and EL35 while the expression of *IAA-amino acid hydrolase 6* decreased in this cultivar at EL32 (Fig. 5). On the other hand, at EL35 levels of IAA seem to be mainly regulated in the two infected cultivars by different expression of *Indole-3-acetic acid-amido synthetase*, suggesting that free IAA levels are regulated at different stages by active mechanisms of hydrolysis and/ or conjugation. Although not statistically significant (p<0,094), the higher basal levels of expression of *IAA-amino acid hydrolase 6* in Syrah grapes at EL32 are accompanied by higher IAA content observed at this stage. Additionally, at EL32 only infected Trincadeira berries showed increased expression of the gene coding for auxin responsive SAUR29 whereas *AUX1* increased in Syrah infected grapes at EL38. This suggests different regulation of auxin signaling and transport during ripening of infected berries from different cultivars.

On the other hand, SA levels were shown to accumulate more upon infection at EL32 in Trincadeira, and at EL35 in Syrah grapes (Fig. 6). At the harvest stage, SA content was relatively the same in both grape cultivars after being infected. Interestingly, the green stage was marked by high levels of expression of the genes coding for EDS1 and PAD4 in infected Trincadeira grapes. Accordingly, Trincadeira grapes showed a larger increase of SA upon infection at this stage. Noteworthy is the higher basal level of expression of *PAD4* in Syrah grapes at EL32 together with higher basal levels of SA. *EDS1* and *PR1* seem to follow the same trend but the differences were not statistically significant. *EDS1* and *PAD4* expression levels were lower at *veraison* and at harvest in both cultivars but *PR1* increased at EL38 especially in infected Trincadeira berries.

In what concerns ABA, it displayed a larger increase in concentration in the Trincadeira cultivar upon infection, in particular at EL32 (Fig. 7). On the other hand, at EL35 infected Syrah berries tended to present higher content in ABA, higher expression of *ABA 8’-hydroxylase* and *ABA receptor PYL4 RCAR10* when compared with infected Trincadeira berries.

The expression pattern of *NCED* at EL32 and EL35 was very similar to the observed content in ABA for the variety of samples but not at EL38. The expression of the gene coding for ABA receptor PYL4 RCAR10 was higher at green stage, especially in Syrah grapes. At EL38, the expression of this gene decreased in Trincadeira berries upon infection. At this stage, infected Trincadeira berries exhibiting heavier symptoms of infection presented higher content in ABA than infected Syrah berries.

## 4 Discussion

Hormones play an important role in plant development and stress responses. Disclosing the roles hormones have in grape ripening and grape defense against major fungal pathogens will enable improvement of fruit traits and productivity. In this context, hormonomics which can be considered as a part of metabolomics, may provide invaluable cues concerning hormonal crosstalk occurring in diverse plant processes. However, in general, concentrations of plant hormones are very low. Most plant hormones cannot be identified in common metabolome analysis therefore a specialized and highly sensitive MS detection system is required to analyze them [52]. Hormonomics has been rarely reported in fruit research, pathogen response and also in general plant science. Oikawa *et al.* [53] have reported comprehensive phytohormone analysis on pear. The majority of plant hormones (e.g. IAA, JA, JA-Ile, SA) presented high content in the youngest fruit and then decreased dramatically. ABA was among the few hormones that increased in the ripening stage. Similar results were obtained in the present study focusing in grape ripening and they were confirmed for two cultivars bringing robustness to the study.

### 4.1 Combined study of two cultivars highlighted differences in the metabolism of ABA, auxins and jasmonates during grape ripening that may influence berry quality

The important role played by ABA in the onset of grape ripening has been widely referred [1,54–56]. Castellarin *et al.* [57] proposed a timeline of events leading to the onset of ripening with increases in ABA occurring early during softening and in the absence of significant increases in expression of the *V. vinifera* 9-cis-epoxycarotenoid dioxygenases. These increases in ABA were accompanied by decreases in a product of ABA catabolism, diphasic acid, suggesting that initial increases in ABA may be due to decreases in catabolism and/or exogenous import. The simultaneous study of grapes from two cultivars collected in the same *terroir* validated the great increase in ABA at initial *veraison* and a putative decrease in ABA catabolism. However, differences between the two cultivars were more pronounced in relation to the expression of *NCED* at EL32 and EL35. Syrah also exhibited a lesser abrupt decrease in ABA catabolism throughout ripening as assessed by the expression of *ABA 8’-hydroxylas*e, which was lower in Trincadeira at EL35 and EL38. Furthermore, the metabolism of ABA is also different between Syrah and Trincadeira at harvest stage with Trincadeira accumulating significantly more ABA (likely due to a higher decrease in ABA catabolism) and transcripts of a gene coding for ABA receptor PYL4 RCAR10 but registering no significant difference in *NCED* expression.

Therefore, ABA biosynthesis, catabolism and signaling are dependent on the cultivar and also on the ripening stage. Recently, the short-term effects of ABA on different organs of grapevine were studied and showed that the responses of each organ were unique indicating that ABA signaling varies with the organ [58,59]. It may be that it also varies with the tissue; therefore differences in ABA metabolism between skin and pulp may partially account for the differences noticed here between the whole berries of the two cultivars. Additionally, ABA may have an impact in fruit quality [60,61] and Trincadeira and Syrah are known to produce grapes and wines with different features. In fact, recent studies have shown that exogenous treatment of pre-*veraison* grape berries with ABA activated the expression of genes involved in cell wall modification, lipid, carbohydrate and flavonoid metabolisms [56].

Contents in auxins and jasmonates are known to decrease during grape ripening while some signaling processes of these growth regulators are activated [1,19]. However, these aspects have not been previously compared between cultivars collected in the same *terroir*. Interestingly, the contents in IAA and the expression of a gene coding for auxin-responsive SAUR29 varied in between the cultivars across the developmental stages. Previously, a role of SAUR proteins in promoting cell expansion was described [62] and this process occurs during berry ripening. Additionally, differences in the expression of gene coding *AUX 1* were noticed between the two cultivars at harvest but not in genes coding for enzymes involved in auxin glycosylation/ hydrolysis though higher content in IAA was observed in Trincadeira at this stage. Since auxin is known to delay increases in berry size, sugar accumulation, and anthocyanin content [13,14,16] the fine regulation of the pool of free IAA and its conjugates together with auxin transport (e.g. AUX1) and signaling may have an impact on final berry quality. Moreover, auxin negatively regulates ABA-induced ripening processes [16] and interactions between ethylene and auxin were reported to be fundamental to the control of berry ripening [63].

Another level of complexity can be added with the participation of jasmonates in the regulation of berry development and ripening as previously suggested [20]. While differences in JA-Ile content were not noticed between cultivars at harvest stage, they were noticed at EL32 and EL35 with Syrah exhibiting more JA-Ile and higher expression of *MYC2* and *jasmonate ZIM domain-containing protein 8 (JAZ8)*. Recently, a putative mechanistic link was reported connecting ABA and JA signaling pathways through a direct interaction of the ABA receptor PYL6 (RCAR9) with the transcription factor MYC2 [51]. The expression of the two genes coding for these receptor and transcription factor followed a similar pattern at EL32 and EL35 for both cultivars though expression levels were generally higher in Syrah. The same was not verified at EL38 when comparing both cultivars indicating that the interplay among hormones is strictly regulated during ripening and depends on the genetic background. In fact, many examples of crosstalk among hormones have been described in grape [1]; the present results suggest that this crosstalk is likely to occur with specificities associated with each cultivar.

Additionally, a new molecular mechanism through which SA antagonizes ABA signaling has been described [64]. Salicylic acid may delay ripening as suggested by the decrease in its content in both cultivars. Additionally, SA treatment has been found to delay the ripening of banana, kiwi and grape fruits [65–67]. Interestingly, PAD4 and EDS1 were shown to regulate cell wall properties in poplar [68] and may eventually play a role in regulation of berry softening. Interestingly, at *veraison* and harvest stage no clear differences were noticed between cultivars in what concerns the content in this growth regulator as well as in the expression of genes coding for *EDS, PAD4* and *PR1*. Though it can be suggested that the role of SA in influencing berry quality characteristics specific of each cultivar may be less determinant than other hormones, its function in grape ripening may still be uncovered.

### 4.2 *Tolerance against* Botrytis cinerea *may be determined by higher basal content in salicylic acid, jasmonates and auxins and additional induced synthesis of salicylic acid*

The analysis of the hormonome showed that Syrah presented at EL32 higher content in SA, JA-Ile and IAA, suggesting that these hormones are involved in basal resistance against *Botrytis cinerea*. Salicylic acid is widely known as determinant for the establishment of basal defenses, effector-triggered immunity, and both local and systemic acquired resistance [69–71]. Additionally, SA has been reported to be involved in the activation of plant defenses against biotrophs and hemibiotrophs, and it also appears to enhance susceptibility to necrotrophs by antagonizing the JA signaling pathway and by inhibition of auxin signaling [23,25,72]. However, SA and JA signaling pathways have also been reported to be either antagonistic or synergistic [72–74]. Our results suggest that SA may also participate in mechanisms associated with resistance/ tolerance to *Botrytis cinerea* since the content in SA was higher in infected green but mainly infected *veraison* Syrah berries comparing to Trincadeira. At this stage only Syrah exhibited mild symptoms due to infection. Previous studies suggest a role of SA in response to infection with *B. cinerea* [75,76]. In tomato, the balance between SA and JA responses seems to be crucial for resistance of unripe fruit to *B. cinerea* [26]. The interaction of JA and SA in promoting resistance has been previously reported in *Arabidopsis* infected with a biotroph and it can depend on the accession [77]. It has been also mentioned that depending on the plant species, the function of SA in immune responses may vary [78,79]. Previously, azelaic acid was identified as a positive marker of infection of Trincadeira green (EL33) and *veraison* (EL35) grapes with *B. cinerea* [4]. This compound is involved in priming the faster and stronger accumulation of salicylic acid in response to pathogen infection in *Arabidopsis* [80]. This accumulation of SA was observed for green Trincadeira grapes but not for *veraison* grapes, highlighting the complex regulation of hormonal metabolism that occurs in response to infection at different stages of ripening. It may be that the pathogen activates the azelaic acid response, but the priming or another downstream process is suppressed or not recognized later during ripening and therefore systemic acquired resistance and the salicylic acid response is not activated as previously suggested [4]. Altogether, it is therefore tempting to speculate that different cultivars may have a different JA and SA balance also in response to necrotrophs and the associated immune response may be therefore distinct. Additionally, only at EL38 Syrah presented significant symptoms of infection and at this stage SA content is similar between the two cultivars.

In basal and in effector-triggered immunity, EDS1 with its direct partner Phytoalexin Deficient4 (PAD4), promotes SA accumulation, and current models position EDS1/PAD4 upstream of SA signaling [50,81]. Recently, an early function of EDS1/PAD4 signaling independent of generated SA has been described [82]. This may justify the significant increase in *EDS1* and *PAD4* expression in infected Trincadeira berries at EL32 without reaching higher content in SA than Syrah. Previously, leaves of the resistant variety Norton infected with powdery mildew, a biotrophic pathogen, presented a constitutively high SA content as compared to the susceptible Cabernet Sauvignon [83]. Additionally, *EDS1* was constitutively expressed to high levels in Norton but not Cabernet Sauvignon, and in Cabernet Sauvignon *EDS1* was induced by PM [84]. Similar results were obtained in this study for *EDS1* and *PAD4* highlighting that some regulatory mechanisms of defense responses may be common to necrotrophic and biotrophic pathogens. On the other hand, EDS1 and PAD4 have been indicated as negative regulators of ethylene/jasmonic acid defense signaling [85], raising the question on whether the high increase in *EDS1* and *PAD4* expression in infected green Trincadeira berries may lead to impairment in defense associated with JA in this cultivar which is a crucial hormone in response to *B. cinerea* [79].

The role played by jasmonates in grape response against *Botrytis cinerea* has been previously referred [4,20]. Interestingly, in tomato transcriptional reprogramming of important JA-signaling components (e.g., *MYC2*) was not evident during fruit infection or during ripening which may indicate that activation of JA-related defenses in fruit occurs via other signaling pathways [26]. However, *MYC2* was up-regulated in infected Syrah grapes at EL35 and EL38 comparing with Trincadeira highlighting the importance of conducting studies in non-vegetative tissues and of comparing climacteric and non-climacteric fruits. In addition, Syrah presented at EL32 and EL35 higher basal levels of expression of *MYC2* than Trincadeira. Mutants of *myc2* have been reported to present reduced resistance to *B. cinerea* [86] indicating that MYC2 may be a positive regulator of defense.

It should be also taken into account that pathogenic fungi can produce plant hormones and manipulate hormonal regulation of plant defense [87]. The secretion of fungal 12-OH-JA can block JA-mediated signaling to suppress the defense response during host penetration [88]. In other cases, pathogen effectors target SA signaling for virulence, by preventing SA accumulation [89]. However, there is no direct evidence that fungal SA or JA is required for their virulence [90].

Regarding auxins, little is known about their role in plant resistance to necrotrophs. However, auxin signaling has been reported to be important for innate immunity, the activation of the auxin pathway mediates pathogen-associated molecular patterns (PAMP)-triggered susceptibility, and auxin opposing regulation mediates PAMP-triggered immunity (reviewed by Naseem *et al.* [91]).

Previously, it was suggested an interaction between auxin and jasmonic acid in resistance to necrotrophic pathogens [32]. Considering the higher basal levels in IAA and JA-Ile in Syrah green berries it may be that these growth regulators are interacting in order to provide a fast response to pathogen attack leading to tolerance of this cultivar.

However, it has also been reported that several plant pathogens can directly synthesize auxin or induce plant auxin biosynthesis or alternatively modulate auxin signaling to render the host more susceptible to infection [29,92–94]. At EL32 infected Trincadeira berries presented increase in IAA content and up-regulation of *auxin-responsive SAUR29* comparing to the control indicating that auxin signaling is activated upon the infection. Additionally, at EL35 infected Trincadeira berries presented higher content in IAA than infected Syrah berries and tendency for higher expression of *AUX1.* This gene was also up-regulated at EL38 in Syrah berries comparing to the control. At this stage, Syrah already presented significant symptoms of infection suggesting that the pathogen may influence auxin transport. Furthermore, the *aux1* mutant, which is defective in auxin influx, cannot develop induced systemic resistance against *Botrytis cinerea* [49]. Our analysis do not seem to corroborate this with increased expression of *AUX1* observed first for infected Trincadeira grapes at EL35 comparing to infected Syrah and an additional increase noted for Syrah at EL38 when they are both exhibiting heavy symptoms of infection. However, *F. oxysporum,* a hemibiotrophic pathogen, requires components of auxin signaling and transport to colonize the plant more effectively, suggesting that alteration of polar auxin transport may also confer increased pathogen resistance [29]. It should also be taken into account that *B. cinerea* can synthesize auxin [95], but the exact function of the pathogen-derived IAA during infection and in interaction with the plant host has not been elucidated.

Salicylic acid has been reported to induce the transcription of genes coding for IAA-conjugating enzyme GH3.5 which converts free IAA into IAA Asp (inactive auxin) [96]. Generally, higher SA levels reduce the pool of active IAA and repress auxin signaling leading to enhanced defense and reciprocally, SA-mediated defenses are attenuated by auxin (reviewed by Naseem *et al.* [91]). This was noticed at EL35 when comparing both cultivars. On the other hand, both cultivars at EL38 exhibited heavy symptoms of infection accompanied by lower levels of SA and increased expression of *auxin-responsive SAUR29* which is in line with previous observations that increased auxin signaling leads to significant reduction in SA accumulation after pathogen infection [96,97].

ABA can also influence the outcome of plant–microbe interactions; in particular the effect of ABA in response against necrotrophic pathogens appears to be complex [79,98–100]. Abuqamar and co-workers [101] suggested a link between ABA, the cell wall, and resistance towards *B. cinerea* in Arabidopsis with ABA treatment inducing the expression of a cell wall loosening gene and contributing to susceptibility. In tomato, increased expression of tomato *NCED* occurs during early infection of susceptible fruit, which suggests a link between ABA synthesis and fruit susceptibility [26]. Our results suggest that ABA is involved in susceptibility of Trincadeira given the high increase in this growth regulator together with increased expression of *NCED* in infected grapes at EL32 comparing to control, though it can be also due to an acceleration of ripening and/ or dehydration promoted by the fungus. This was not noticed in Syrah at this stage; instead Syrah present higher basal expression of *ABA receptor PYL4 RCAR10,* so ABA signaling processes can eventually be connected to basal resistance in interaction with other growth regulators such as JA and SA [102,103]. In fact, negative and positive roles have been described for this hormone depending on the pathosystem, developmental stage of the host, and/or the environmental conditions in which the plant–pathogen interaction occurs [33,104,105]. The results presented here suggest that it may also depend on intra-species genetic variation of hormone networks (Fig. 8).

**Figure 8:**
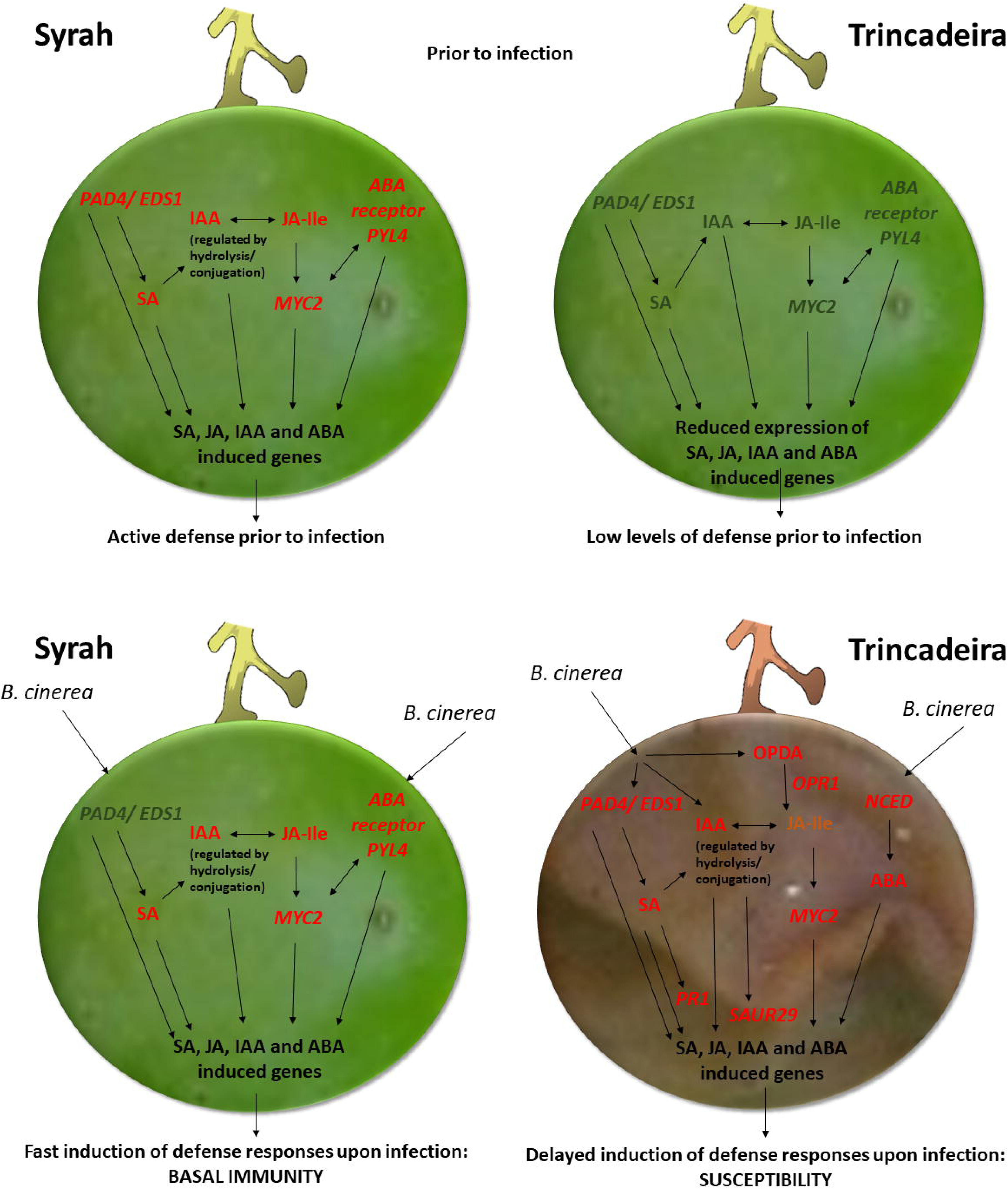
Hormonal metabolism in grapes from green clusters (EL32) before and after infection with *B. cinerea* and representing tolerant (Syrah) and susceptible (Trincadeira) genotypes. High levels are highlighted in red and low levels in green. Basal levels of hormones (IAA, SA and JA-Ile) in Syrah are putatively involved in a faster response to *Botrytis cinerea.* Jasmonates, ABA and SA signaling are also activated in Syrah grapes without infection. Trincadeira presents at green stage a more extensive reprogramming of hormonal metabolism upon infection (high increase in ABA, IAA, SA and JA-Ile content together with activation of jasmonate, IAA and SA signaling), but this is likely to result in delayed defense responses and inefficient defense. The kinetics of phytohormone biosynthesis and the timing and sequence of initiation of hormones` signaling during the interaction of fruit with *B. cinerea* may determine the outcome of hormones’ interaction and successful defense.

## 5 Conclusions

The combined analysis of the hormonal profile and targeted qPCR analysis of genes involved in hormonal metabolism during ripening and upon *B. cinerea* infection suggested new roles for SA, IAA and ABA in cultivar fruit specificities and in response to necrotrophic pathogens. High basal levels of SA and IAA at an early stage of ripening, together with activated SA and IAA metabolism and signaling (Fig. 8), and possibly in interaction with JA, seem to be important in providing a fast defense response leading to grape tolerance against *B. cinerea*.

The results indicate that the role of plant hormones in promoting fruit resistance or susceptibility is extremely complex depending not only on the relative content of the hormones, but also on the timing of the synthesis and perception of the hormones. Additionally, other factors may be important as well, namely the competence of the host tissue to respond to active forms of the hormones, the localization of the plant’s response to the hormones, the pathogen’s infection strategy, including its own production of hormones [26,69]. Moreover, the balance and interaction among hormones may be fundamental in providing either resistance or susceptibility; two genetic backgrounds in the same plant species may present different hormone signaling requirements for resistance[31] as indicated here. Further studies on inter- and intraspecies genetic variation of hormone networks may validate the importance of basal hormonal levels in resistance/ tolerance as suggested here and might provide insights into possible strategies of manipulation using genome-editing technologies. In grapevine, the limitation will be the optimization of transgenesis’ protocols for different cultivars enabling functional gene characterization in grapes [106]. Additionally, further hormonomics and RNA sequencing applied to the specific study of cells and tissues will elucidate unknown functions of plant hormones in fruit growth, development and quality as well as in response against pathogens.

## Author contributions

J.C., M. A-Trapp, D.P., F.S. and AMF carried out the experiments; P.R and C.R. performed the infections. AMF designed the study and wrote the manuscript with input from A.M.

## Supporting information

## Acknowledgments

This work was supported by the Portuguese Fundação para a Ciência e Tecnologia (FCT) [grant numbers UID/Multi/04046/2013, FCT Investigator IF/00169/2015, Fellowships PD/BD/114385/2016 and PB/BD/130976/2017]. Ana M. Fortes and Marilia Almeida-Trapp gratefully acknowledge financial support by the Innovation Prize CNOIV 2016 and by a Capes-Humboldt Research Fellowship, respectively.

## Appendix. Supplementary data

Supplementary Table S1-List of primers used in real-time reverse transcription-PCR. Primers were designed using Primer Select from DNAstar package.

Supplementary Figure S1-Hormone Metabolic Pathways and Genes. A, The enzyme AOS catalyzes a reaction of the jasmonic acid pathway, leading to the formation of the precursor OPDA; B, The enzyme OPR1 catalyzes a step in the jasmonic acid/ jasmonoyl-isoleucine pathway in which OPDA is precursor; C, In the absence of JA-Ile, JAZ8 acts as a repressor of jasmonate signaling by inhibiting the activity of MYC2. When present, JA-Ile promotes the formation of a JAZ8 complex, allowing the released MYC2 to promote the expression of its target genes; D, The enzyme 9-cis-epoxycarotenoid dioxygenase (NCED) catalyzes a reaction in the abscisic acid pathway; E, The presence of ABA activates the inhibition of PP2C phosphatases by the ABA receptor complex, thus promoting the transcription of ABA-responsive genes; F, By intervening in abscisic acid catabolism, the enzyme ABA 8’-hydroxylase plays a regulatory role in controlling the level of ABA in plants. The product of ABA 8’-hydroxylase is 8’-hydroxy-abscisic acid; G, IAA-aminoacid synthetases conjugate IAA with target aminoacids as a way to regulate the pool of active auxin. IAA-aminoacid hydrolases act as a counterpart to the IAA inactivating function of the synthetases; H, Aux1 is a carrier protein involved in the influx of auxin to the interior of the cell. When auxin is present, the expression of auxin-responsive genes, including the auxin-responsive SAUR29, is activated; I, EDS1 operates upstream of SA-dependent defenses, promoting the establishment of the hypersensitive response and effector-triggered immunity. Another function of EDS1 involves the recruitment of PAD4, potentiating basal plant defenses through the transcription of defense-related genes and further accumulation of SA. The expression of both *EDS1* and *PAD4* has been described as regulated by SA through a positive feedback loop.

