## Supplementary figures and images for "The study of hormonal metabolism of Trincadeira and Syrah cultivars indicates new roles of salicylic acid, jasmonates, ABA and IAA during grape ripening and upon infection with *Botrytis cinerea*"

### Supplementary file 1

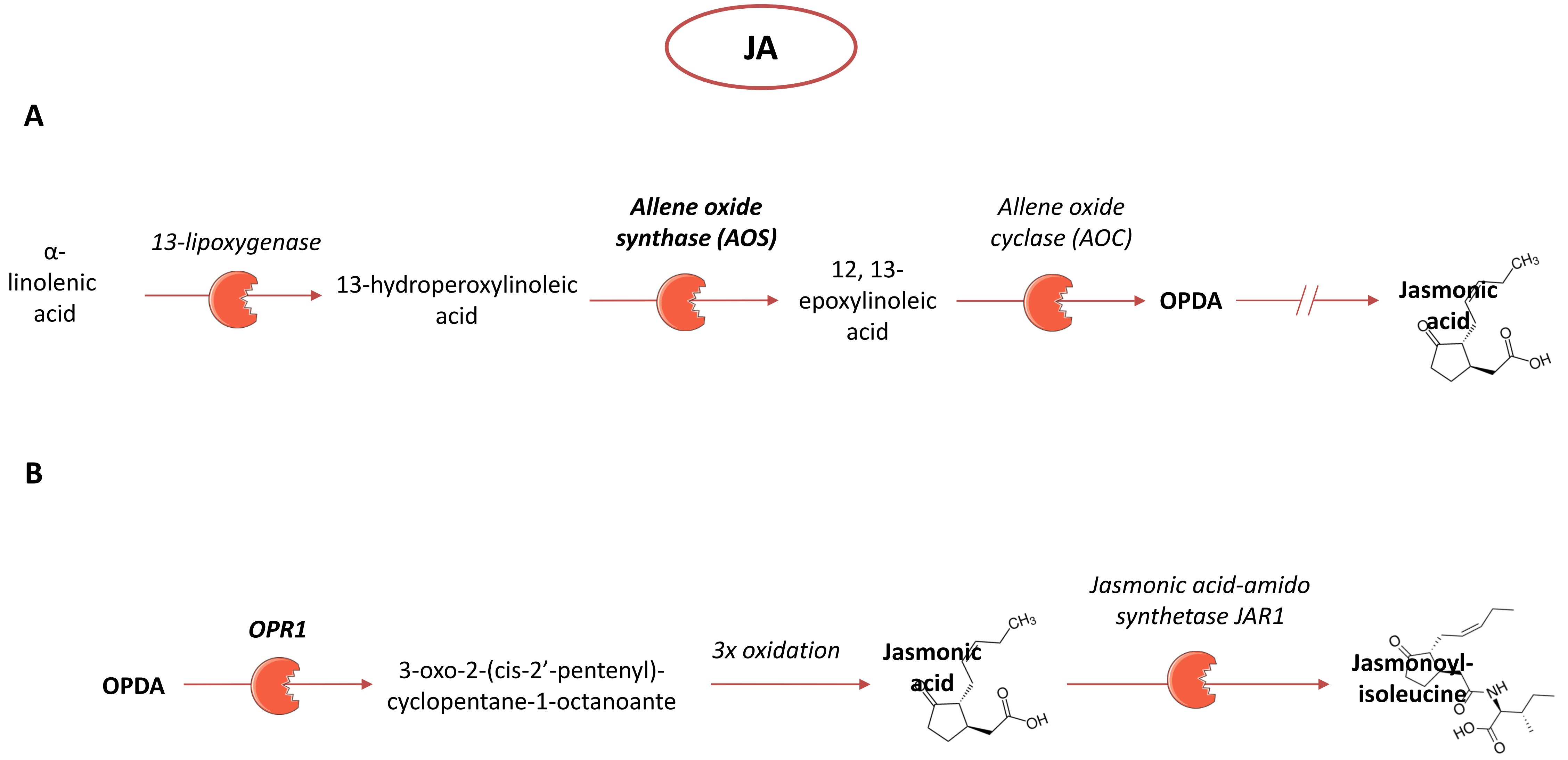

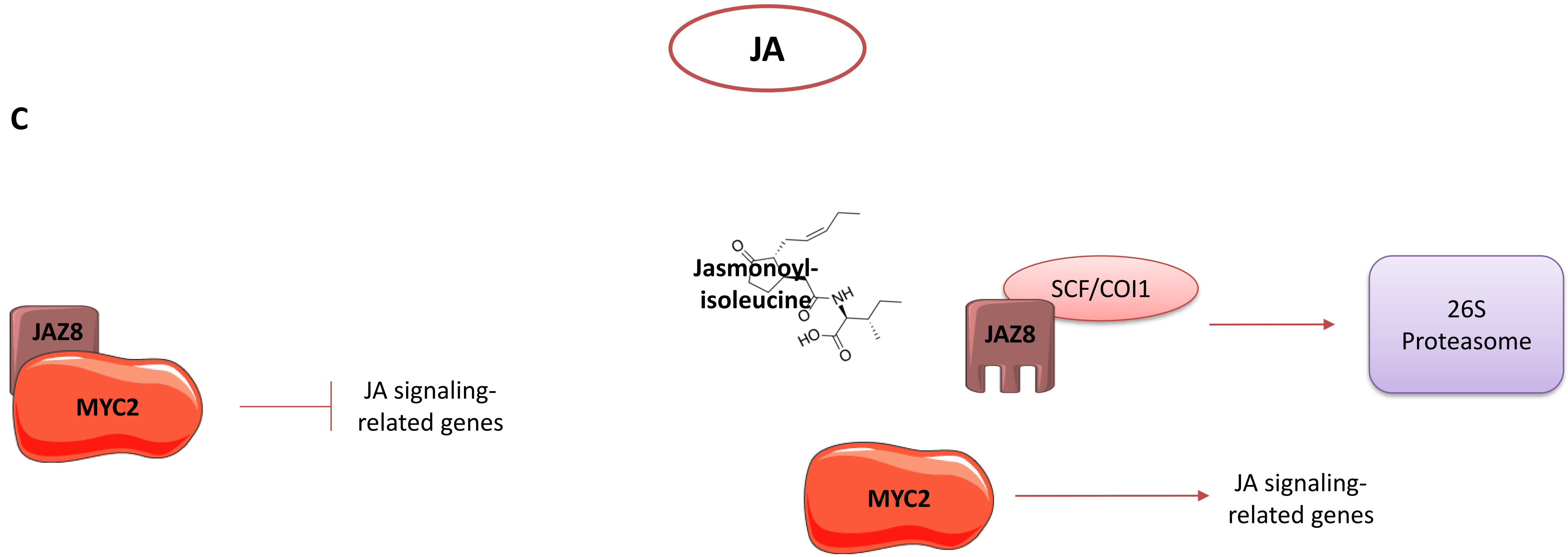

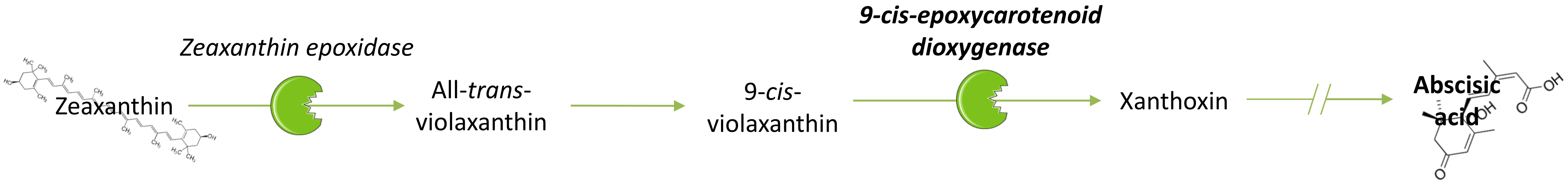

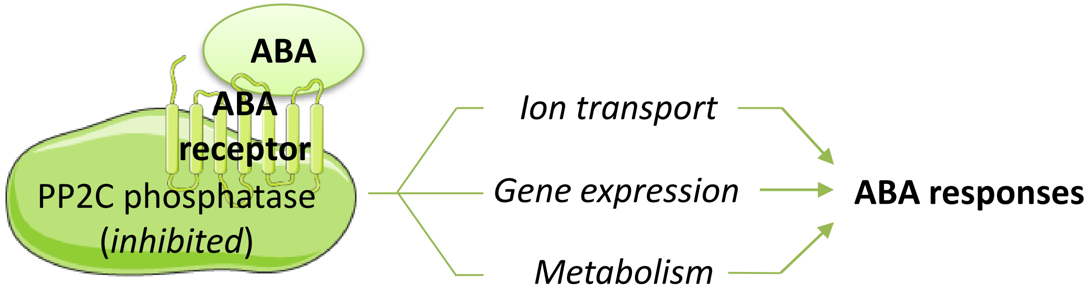

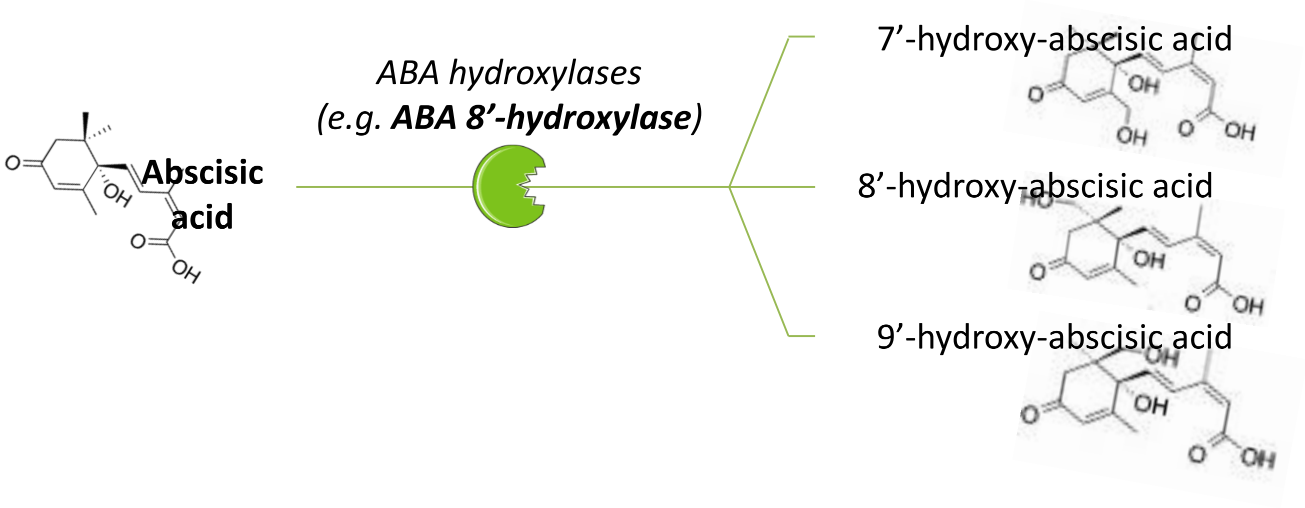

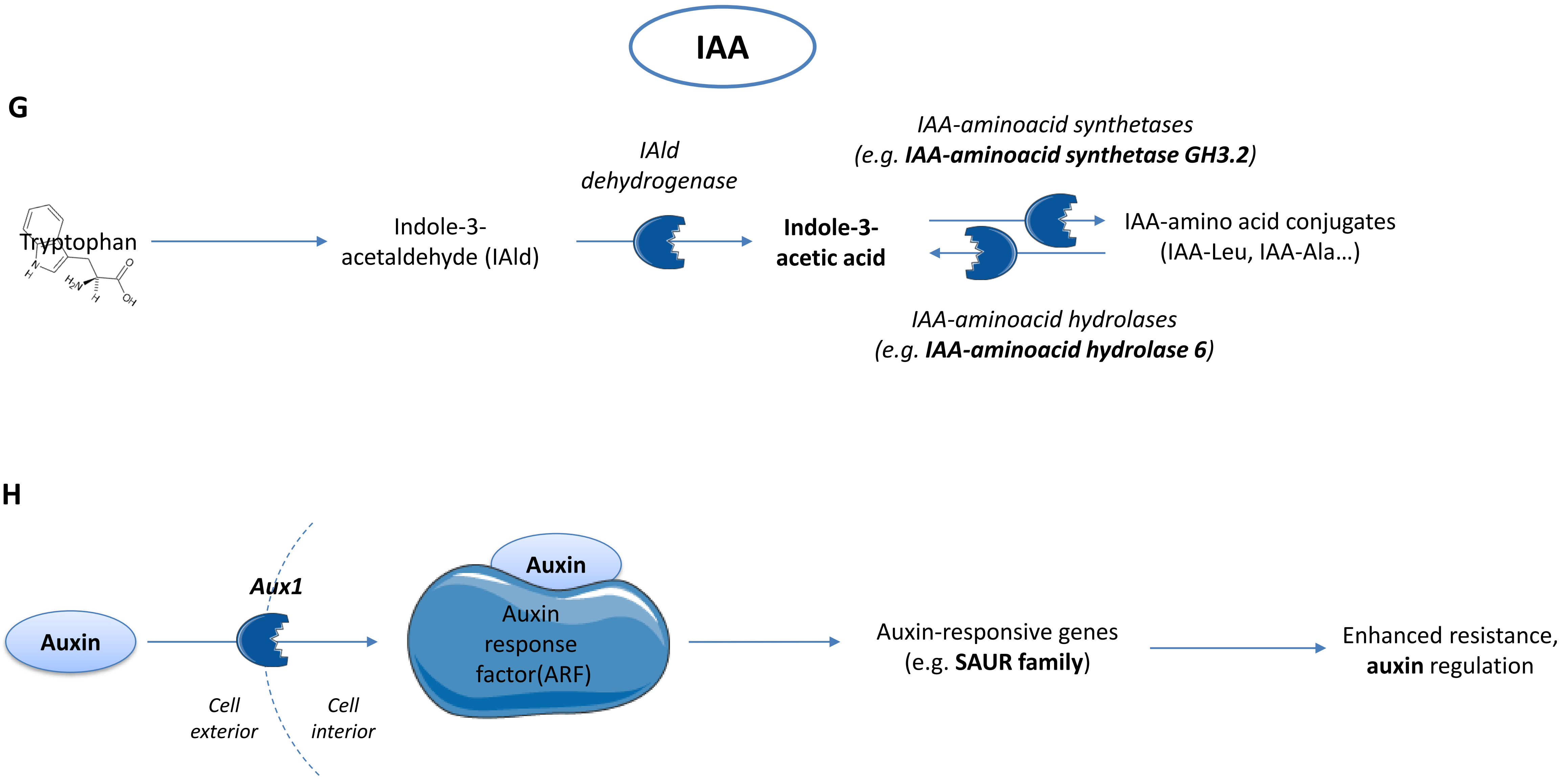

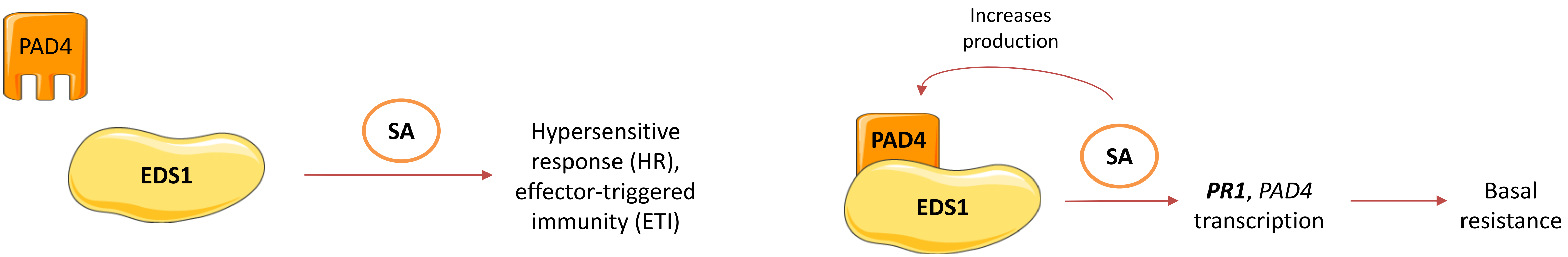
