## Supplementary material for "The study of hormonal metabolism of Trincadeira and Syrah cultivars indicates new roles of salicylic acid, jasmonates, ABA and IAA during grape ripening and upon infection with *Botrytis cinerea*"

| Target sequences and Probe ID |  | Primer sequence | T_melting_  (°C) |
| --- | --- | --- | --- |
| Elongation Factor 1 α (EF1α)  *VIT_06s0004g03220* | Fw Rev | CGTCATAGTTTTCTGCCTTCTTCC  TGCCACCGCCTATCAAGC | 59.3  59.3 |
| Actin  *VIT_04s0044g00580* | Fw Rev | GGTCAACCATGTTCCCTGGTATT  GGAGCAAGAGCAGTGATTTCCTT | 59.8  59.9 |
| Allene oxide synthase (AOS)  *VIT_03s0063g01820* | Fw  Rev | GCTTTACCGCGCCTTTTATGC  TCCTGCTGAGCCAACCCACTT | 57,7  58,3 |
| 12-oxophytodienoate reductase 1 (OPR1)  *VIT_18s0041g02070* | Fw  Rev | CCCCGGGTATATGGACAAAAGA  CCACATGCCAAAGCTGACAAAT | 56,6  56,4 |
| Jasmonate ZIM domain-containing protein 8 (JAZ8)  *VIT_210s0003g03790* | Fw  Rev | CGGAAGGATCTGCGTTTGTGA  TGAGGCAGTCGGGGTTCTTGT | 57,4  58,2 |
| Transcription factor myc2-like (MYC2)  *VIT_202s0012g01320* | Fw  Rev | CAGTGGTTCGGGAGGCAGATAG  CACAGCTCGGAGGGCATAAAAC | 57,4  57,5 |
| 9-*cis*-epoxycarotenoid dioxygenase (NCED)  *VIT_10s0003g03750* | Fw  Rev | CGCTCGCCTCCTCCTCTTCTAT  AGGGCTTGATTCGCACTTGGTA | 57,9  57,5 |
| ABA receptor PYL4 RCAR10  *VIT_08s0058g00470* | Fw  Rev | TGCCGCCGCGAATAACCATA  GACGGCGGAGCAGCATTGATT | 60,7  60,9 |
| ABA 8’-hydroxilase  *VIT_18s0001g10500* | Fw  Rev | CAAGCCCACATACCCCAAAAGT  TCAATATCGGCCACCAAGTTCC | 56,7  57,5 |
| IAA-amido synthetase GH3.2  *VIT_07s0129g00660* | Fw  Rev | GAGGCCATTCTTTGCGTTGACT  CGACTCGGAGGACTTCTTTGTG | 57,1  55,4 |
| Auxin-responsive SAUR29  *VIT_16s0098g01150* | Fw  Rev | GGGGAGGAGCAGCAGAGGTTTG  GGCTGTGGTGGTGGTGGGACTT | 60,9  61,7 |
| IAA aminoacid hydrolase 6  *VIT_18s0001g02570* | Fw  Rev | ACTCCAAGGCTCCAACCACTCA  TTTCAACCACTCCACCGTCTCC | 57,3  57,6 |
| Auxin transporter protein 1 (Aux1)  *VIT_213s0067g00330* | Fw  Rev | TGGTGGGGGTGCATGATACAAA  ATGGGCAAGAGCTGGGATGATG | 59,4  59,8 |
| Enhanced disease susceptibility 1 (EDS1)  *VIT_17s0000g07420* | Fw  Rev | GAGCTTCCGGTGTCTTCTGATG  TTTCGCTTCTCCAACTCTCCTG | 58,1  60,1 |
| Phytoalexine deficient 4 (PAD4)  *VIT_07s0031g02390* | Fw  Rev | GGCTAGCTGGGCAGGAGTCAA  AGGTGTGGCGGTAACGGATTCA | 55,3  55,2 |
| Pathogenesis-related protein 1 (pr1)  *VIT_03s0088g00700* | Fw  Rev | TGCCTACGCCCAGAACTATGC  TGCCTGTCAATGAACCACTGC | 56,7  55,8 |
